# SAMP V2: A novel stacking ensemble learning model for antimicrobial peptides identification based on augmented split amino acid composition with biochemical-sequence-order information

**DOI:** 10.64898/2026.08.12.744552

**Authors:** Mengtao Sun, Jieqiong Wang, Shibiao Wan

## Abstract

Antimicrobial resistance reduces the effectiveness of conventional antibiotics and has become a major global health threat, highlighting the need for new anti-infective agents. Antimicrobial peptides (AMPs), a diverse class of innate immune effectors with broad-spectrum antimicrobial activity, are promising candidates for combating drug-resistant infections. Identifying AMPs by wet-lab experiments, however, remains costly and time-consuming, creating a strong demand for computational identification methods. Our recently developed method, SAMP, captures region-specific residue distributions based on proportionalized split amino acid composition. However, SAMP might ignore key biochemical information and sequence order information. Here we present SAMP V2, a stacking ensemble learning framework based on biochemical and sequence-order information augmented split amino acid composition (BIA-SAAC), which extends SAMP by integrating pseudo-amino acid composition features with biochemical and sequence-order information into split peptide regions. Specifically, each peptide is divided into N-terminal, middle, and C-terminal regions, and pseudo amino acid composition is calculated within each region. Benchmarking tests on six independent test datasets, SAMP V2 outperformed multiple state-of-the-art models, including AMPpred-MFA and iAMP-Attenpred, in terms of accuracy, MCC, G-measure and F1-score. Given its high and robust performance, SAMP V2 could significantly accelerate the discovery of next-generation antimicrobial therapeutics for addressing the global threat of multidrug-resistant pathogens.

## Introduction

The accelerating emergence of antimicrobial resistance has become one of the most severe threats to global public health, undermining decades of progress in infectious disease treatment ^1^. The extensive and often indiscriminate use of antibiotics in clinical medicine, agriculture, and animal husbandry has driven the evolution of multidrug-resistant pathogens ^2–6^, while the discovery of new antibiotic classes has stagnated over recent decades. As resistance continues to erode the efficacy of last-resort antibiotics, there is an urgent demand for alternative antimicrobial strategies that operate through mechanisms distinct from conventional small-molecule drugs ^7^. In this context, antimicrobial peptides (AMPs) have attracted increasing attention as a promising class of next-generation therapeutics. Antimicrobial peptides are short, gene-encoded amino acid sequences that are widely distributed across all domains of life and constitute a central component of innate immunity ^8–12^. Unlike traditional antibiotics that typically target specific intracellular enzymes or metabolic pathways, AMPs predominantly exert their antimicrobial activity through interactions with microbial membranes ^13^. Their amphipathic and cationic nature enables selective binding to negatively charged bacterial membranes, leading to membrane disruption and cell death. This physical mode of action confers AMPs with broad-spectrum efficacy against bacteria, fungi, viruses, and parasites, while simultaneously imposing a high evolutionary barrier to resistance development ^14–18^. In addition to their direct antimicrobial activity, many AMPs exhibit immunomodulatory, anti-inflammatory, and wound-healing functions, further highlighting their therapeutic potential ^19–21^.

Despite their considerable promise, the identification of novel AMPs remains a major bottleneck. Conventional experimental discovery pipelines are time-consuming, costly, and inherently limited in throughput ^22^. Moreover, the combinatorial scale of peptide sequence space far exceeds the capacity of experimental screening ^23^, and advances in genomics and metagenomics have revealed an immense reservoir of uncharacterized peptide sequences ^24^. As a result, experimental approaches alone are limited in their ability to rapidly explore and prioritize candidate AMPs for validation. To overcome these limitations, computational identification of AMPs has emerged as a critical complementary strategy. In silico models enable high-throughput screening of vast peptide libraries and genomic datasets, dramatically narrowing the search space for experimental testing. Over the past decade, a wide range of machine learning (ML)/deep learning (DL) approaches have been developed for AMP identification, progressively incorporating richer representations of peptide composition, sequence order, regional distribution, and structural potential. These computational frameworks not only accelerate AMP discovery but also provide mechanistic insights into the physicochemical determinants underlying antimicrobial activity. In 2018, Bhadra et al. established a foundation with AmPEP, employing random forest ^25^ algorithm to decipher distribution patterns of amino acid physicochemical properties ^26^. Building on sequence-based approaches, Ma et al. (2022) ^27^ adapted natural language processing architectures, integrating long short-term memory (LSTM) ^28^, attention mechanisms, and bidirectional encoder representations from transformers (BERT) ^29^ to successfully mine functional peptides from complex human gut metagenomes. Extending these DL-based strategies, Xing et al. (2024) developed ^30^ iAMP-Attenpred, which leveraged BERT to encode amino acid sequences as contextual representations and integrated convolutional neural network (CNN), bidirectional LSTM, and attention modules to capture local motifs and long-range dependencies, thereby enhancing AMP classification performance across benchmark datasets. Subsequent studies have focused on enhancing predictive robustness through ensemble learning. For instance, Wang et al. (2024) developed E-CLEAP ^31^, which aggregated multilayer perceptron (MLP) classifiers via soft voting to address data imbalance. Li et al. (2024) ^32^ proposed AMPpred-MFA which integrates fragment-level k-mer embeddings with dipeptide deviation from expected mean features through a stacking architecture with CNN, BiLSTM, and multihead attention. Most recently, Sun et al. (2025) ^33^ advanced the field into the structural dimension with the SSFGM-Model, a multimodal geometric deep learning framework that leverages AlphaFold2-predicted structures and graph neural networks ^34^ to interpret physicochemical surface features alongside sequence data.

Additionally, to address the limitation that many existing AMP predictors rely mainly on global compositional and may overlook position-dependent residue patterns, we previously proposed SAMP ^35^, an AMP prediction framework based on proportionalized split amino acid composition (PSAAC). By splitting each peptide into N-terminal, middle, and C-terminal regions, SAMP captured region-specific residue distributions, including terminal patterns that may be ignored when only whole-sequence features are used. SAMP further employed a random projection-based ensemble learning strategy ^36^, in which multiple random projections were used to generate low-dimensional feature spaces and the resulting base classifiers were integrated to improve robustness and scalability for AMP identification. SAMP achieved strong performance across multiple benchmark datasets and outperformed several state-of-the-art AMP prediction tools. However, SAMP primarily focused on regional amino acid composition and did not explicitly model residue-order dependencies or biochemical-order relationships along peptide sequences. In addition, its ensemble architecture relied on homogeneous support vector machine models, which limited model diversity and the ability to capture complementary decision patterns.

To address these concerns, we present SAMP V2, which further extends AMP representation by incorporating order-aware sequence information to better capture the positional and contextual relationships among residues. Compared with the original SAMP, which employed an ensemble random projection strategy combined with PSAAC, SAMP V2 introduces ensemble stacking learning architecture combined with biochemical-information augmented split amino acid composition (BIA-SAAC) which incorporates regional biochemical properties and sequence-order correlations. However, instead of relying solely on amino acid frequency statistics, BIA-SAAC computes pseudo amino acid composition ^37^ within each segment, thereby encoding both local sequence-order information and physicochemical correlation patterns that are essential for antimicrobial activity. To effectively integrate BIA-SAAC with conventional sequence-derived features, SAMP V2 adopts a stacking ensemble framework in which multiple heterogeneous base classifiers are trained independently, and their prediction outputs are subsequently combined by a meta-learner to produce the final decision. This stacked architecture enables SAMP V2 to leverage complementary decision boundaries across different learners, improving robustness and generalization while providing a unified framework for integrating richer, biologically informed feature representations. Additinally, extensive benchmarking experiments demonstrate that SAMP V2 consistently outperforms existing state-of-the-art AMP prediction methods, including SAMP ^35^, amPEPpy ^38^, AMPScanner V2 ^39^, AMPpred-MFA, iAMP-Attenpred and five independent traditional basic classifiers, across multiple evaluation metrics, such as accuracy, the geometric mean of recall and precision (G-measure), Matthews correlation coefficient (MCC), and F1-score. To enhance the impact of SAMP V2, we have developed a software package that is available on our lab’s GitHub repository at https://github.com/wan-mlab/SAMP-V2.

## Materials and methods

### Dataset

To ensure robust model training and evaluation, we curated several datasets from established sources ^10,30,40,41^. A preprocessing pipeline was applied to all raw sequences to ensure data quality: (1) <u>Length filtering</u>: sequences were restricted to a length of 10-500 amino acids; (2) <u>Residue</u> <u>standardization</u>: sequences containing non-canonical amino acids were excluded to strictly retain the 20 standard residues; and (3) <u>Deduplication</u>: redundant sequences among all datasets were removed to prevent data leakage. Following these steps, training dataset was constructed, including 2,021 AMPs and 2,019 non-AMPs sourced from the AMPScanner V2 web server (www.ampscanner.com), incorporating updated APD3 ^10^ sequences. Additionally, three benchmark datasets were constructed and served as the independent test set for model performance evaluation. Benchmark Dataset 1 (benchmark1) collected 572 AMPs and 1920 non-AMPs from iAMP-Attenpred ^30^. Benchmark Dataset 2 (benchmark2) included 4564 AMPs and 5296 non-AMPs sourced from Yan et al ^40^. Benchmark Dataset 3 included amphibian, bacteria, human and plant sourced AMPs and non-AMPs from dbAMP database version 1 ^41^. There were 517 AMP and 932 non-AMP sequences in amphibian dataset, 226 AMP and 4721 non-AMP sequences in bacteria dataset, 39 AMP and 894 non-AMP sequences in human dataset, and 307 AMP and 3185 non-AMP sequences in plant dataset. Dataset composition and sequence-length distributions were compared across all AMP and non-AMP datasets. AMP sequences were mainly derived from the benchmark2 and training datasets, which together accounted for the majority of AMP samples, whereas the non-AMP sequences were mainly derived from benchmark2 and bacteria datasets (Supplementary **Fig. S1A-B**). Density plots further showed heterogeneous length distributions of AMP sequences and non-AMP sequences across datasets (Supplementary **Fig. S1C-D**).

### Feature extraction

To encode peptide sequences into numerical vectors, a comprehensive set of sequence-derived features was extracted by the *Peptide* ^42^ and *protr* ^43^ packages in R, following widely adopted practices in AMP prediction studies. These features were designed to capture both global compositional characteristics and physicochemical properties of peptide sequences. Specifically, three categories of features were considered: (i) amino acid composition (AAC), (ii) structural (STRL) features, including pseudo amino acid composition (PseAAC) and BIA-SAAC, and (iii) physicochemical (PHYC) features, including hydrophobicity, net charge, and isoelectric point (**Table 1**).

**Table 1.**
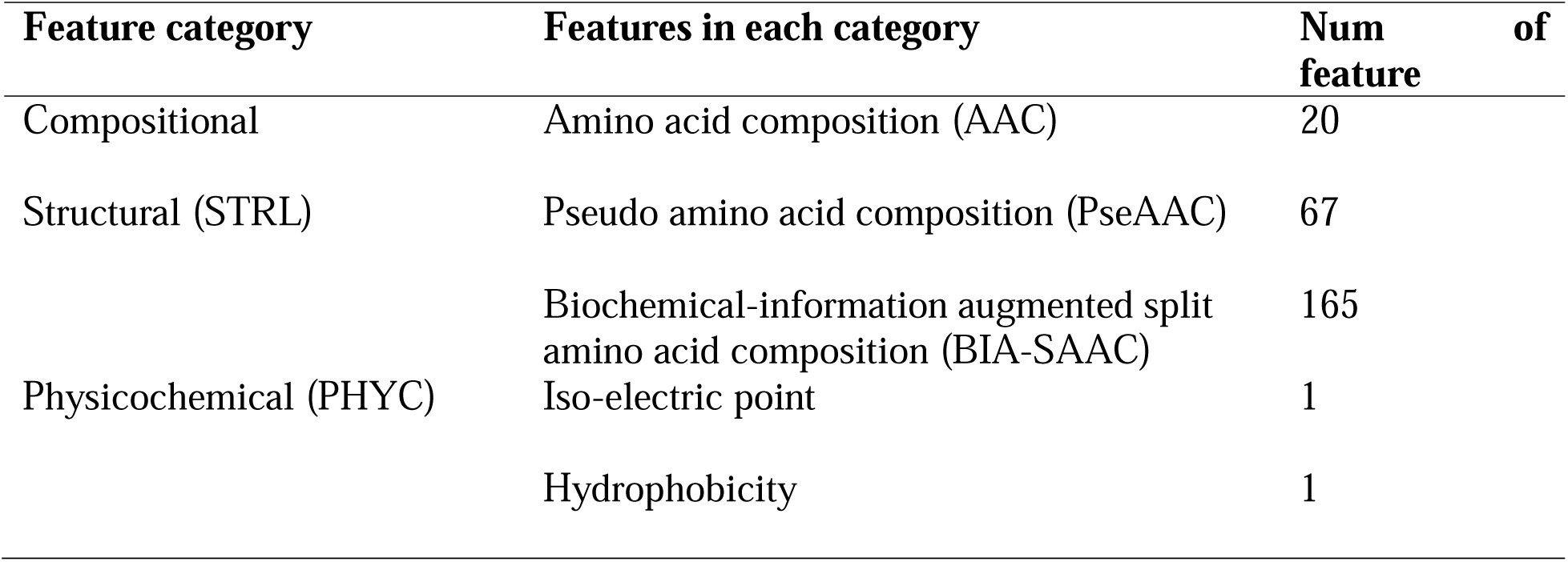
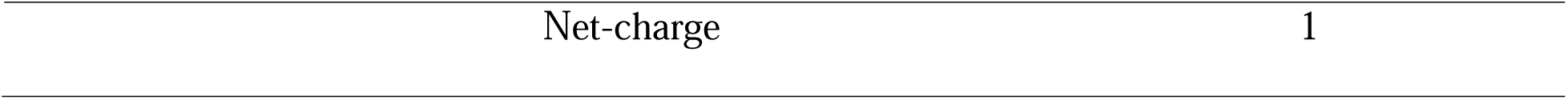
Summary of the feature sets. Compositional, structural and physicochemical features were extracted from sequences, including amino acid composition, pseudo amino acid composition, biochemical-information augmented split amino acid composition, iso-electric point, hydrophobicity, and net-charge, for model training and evaluation. The dimension of PseAAC and BIA-SAAC is based on the lambda=5.

Conventional features including amino acid composition and physicochemical properties. AAC encodes the relative frequency of the 20 standard amino acids, providing a global summary of residue usage while accounting for sequence-length variation. PHYC features were computed to reflect key biophysical properties governing peptide-membrane interactions. Additionally, BIA-SAAC extends simple composition by incorporating biomedical-order correlation information, thereby capturing local physicochemical dependencies between neighboring residues that are important for secondary structure formation and antimicrobial activity.

### The SAMP V2 framework

SAMP V2 is an ensemble stacking-based framework for AMP identification that integrates complementary information across multiple learning stages. Peptide sequences are first encoded into numeric feature representations, including the newly proposed augmented split amino acid composition (BIA-SAAC) (**Fig. 1A**) and conventional sequence-derived features (**Fig. 1B**). At the first level of the stacking architecture, heterogeneous base classifiers are trained independently on the same feature space to capture diverse decision patterns and generate prediction score matrices. At the second level, the base-level prediction scores are concatenated with the original feature matrix and fed into a meta-learner to produce the final classification (**Fig. 1C**).

**Fig. 1.**
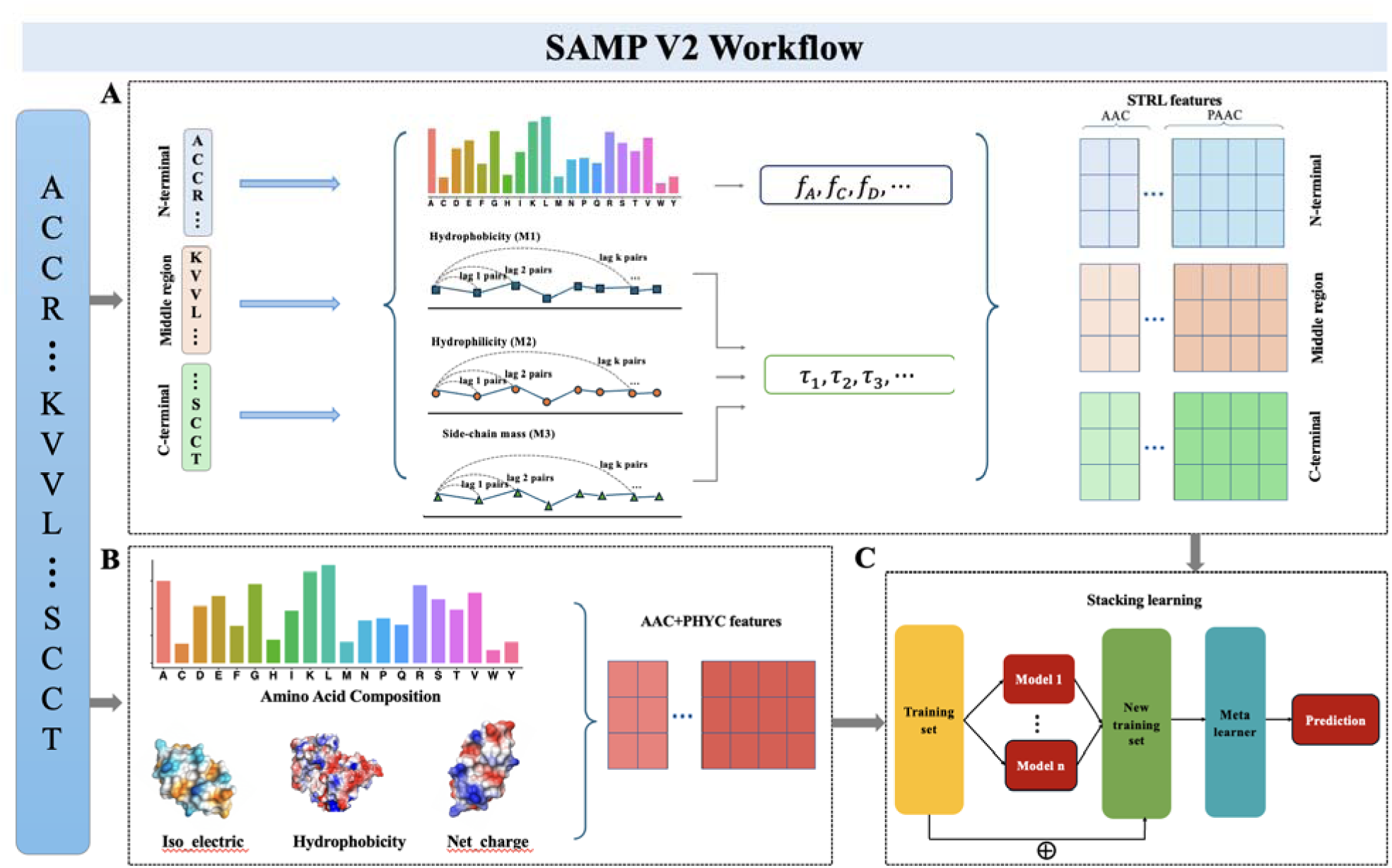
The framework of SAMP V2 based on composition and biochemical information for AMP prediction. Split amino acid sequence into N-terminal, C-terminal and middle regions. For each segment, pseudo amino acid composition is computed to reflect biochemical-sequence-order correlation. The resulting segment-level feature vectors are concatenated to form the final BIA-SAAC feature vector used for downstream classification (**A**). For the residue-level features, we extract four different features categories, including amino acid composition, iso-electric point, hydrophobicity, and net-charge (**B**). The computational pipeline of the SAMP V2 for AMP prediction. Peptide sequences are encoded into numeric features, including conventional descriptors and the proposed BIA-SAAC representation. The feature matrix is fed into base classifiers to generate prediction score matrices. These scores are concatenated with the original features and used as input to a meta-learner, which produces the final AMP prediction (**C**).

In addition to the stacking architecture, BIA-SAAC extends conventional composition-based representations by jointly encoding regional sequence heterogeneity and sequence-order correlations through proportional splitting of peptide sequences into N-terminal, middle, and C-terminal regions. This biologically informed representation enables SAMP V2 to more effectively capture amphiphilicity-related and structural patterns of AMPs. The performance of SAMP V2 was evaluated on independent test datasets and compared with existing state-of-the-art methods, including SAMP, AmPEPpy, and AMPScanner V2, as well as five independently base classifiers trained with identical dataset.

### Biochedical-information augmented split amino acid composition (BIA-SAAC)

To capture region-specific biochemical-sequence-order information while maintaining comparability across peptides of different lengths, we propose BIA-SAAC (**Fig. 1A**). Given a peptide sequence of the length *L*, we split it into three segments (N-terminal, middle, and C-terminal) using user-defined proportions with. The corresponding segment lengths are
(1)

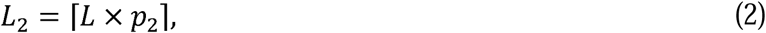

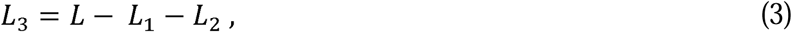

where ‘⌈⌉’ rounds a value up to the nearest integer. The segments are

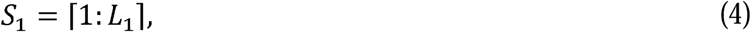

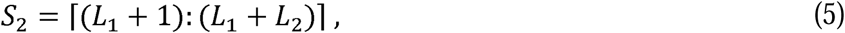

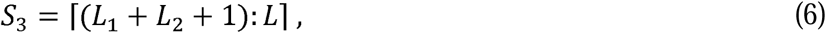

denote the normalized occurrence frequency of the -th amino acid (1,…,20) in For each segment *s_i_* (*i* = 1, 2, 3), we compute its PseAAC vector following Chou’s formulation^37^. Let *f^(i)^_u_* denote the normalized occurrence frequency of the *u*-th(*u*=1,…,20) in segment *s_j_*, and let *τ^(i)^_k_(k = 1, …, λ)* be the k-th tier sequence order correlation factor defined as

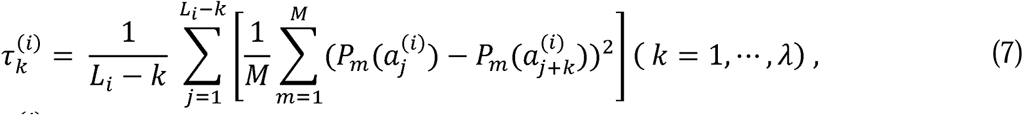

where *a^(i)^_j_* and *L_i_* denote the *j*-th residue and length of segment *S_i_*, espectively.*P_m_* denotes the tier sequence-order correlation within segment by averaging squared physicochemical *m*-th physicochemical property *M* is the number of properties used, and *τ^(i)^_k_* quantifies the *k*-th tier sequence-order correlation within segment *S_i_* by averaging sqaured physicochemical differences between residue pairs *j* and *j + k* (e.g., based on standardized hydrophobicity, hydrophilicity, and side-chain mass).

Using a weight factor *ω* = 0.05, the segment-level PseAAC components are:

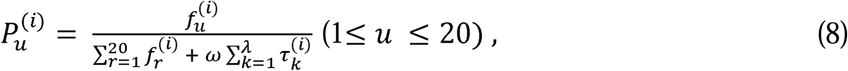

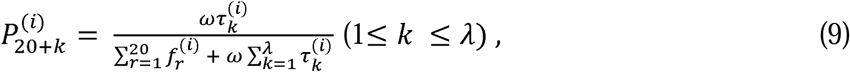

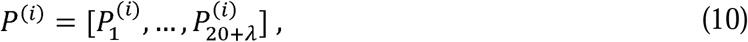

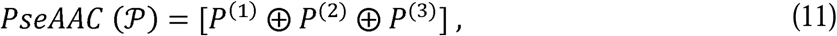

Where ⊕ denotes concatenation.

For each segment *s_i_* (*i* = 1, 2, 3,), we futher its amphiphilic pseudo amino acid composition (APAAC) vector following Chou’s formulation ^37^. Let *M* denote the number of denote the normalized value of the -th physicochemical property for the -th amino acid in physicochemical properties used (e.g., hydrophobicity and hydrophilicity) and let *P_m_(a^(i)^_j_* denote the normalized value of the *m*-th physicochemical property for the *j*-th amino acid in segment *S_i_*. The *k*-th tier sequence-order correlation factor is defined as

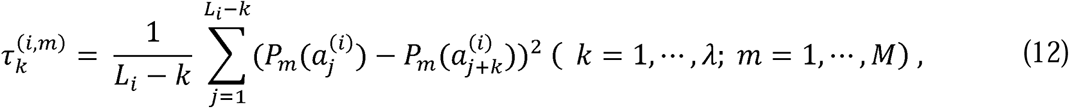

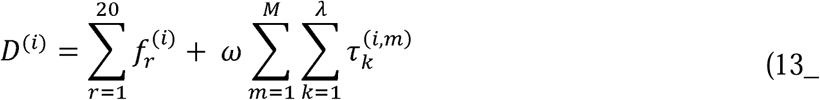

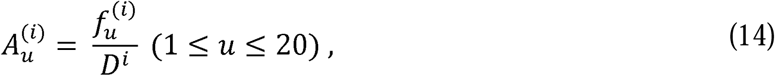

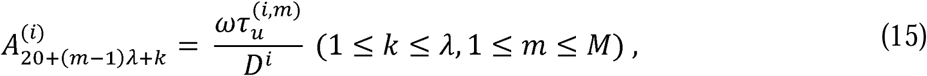

thus, the APAAC vector for segment *S_i_* is

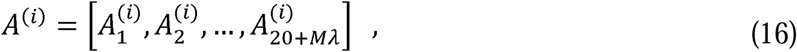

Finally, the BIA-SAAC is obtained by concatenating the segment level PseAAC and APAAC vectors:

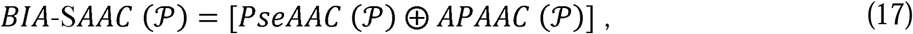

this design preserves segment-specific composition and sequence-order patterns, facilitating robust learning across variable-length peptides.

### Ensemble stacking learning architecture

An ensemble stacking learning framework was employed to integrate heterogeneous classifiers for antimicrobial peptide (AMP) identification (**Fig. 1C**). Multiple sequence-derived features were extracted for each peptide, resulting in a feature matrix **X** ∈ **ℝ***^n^*^×*d*^, where *n* denotes the number of samples and *d* represents the dimensionality of the feature space At the base level, two complementary classifers, *C*_1_ and *C*_2_, were trained independently using **X** as input For each sample both classifers produced class-wise prediction scores specifically, the *C*_1_ and *C*_2_ yielded score matrics

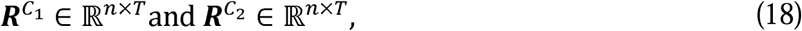

respectively, where *T* is the number of classes.

To construct the meta-level representation, the original feature matrix and the base-level score matrices were concatenated along the feature dimension

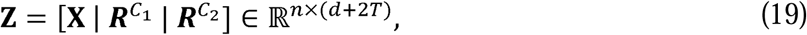

where **Z** is the final score matrix for meta learner training. The base classifier outputs were then concatenated with the original feature matrix to construct an augmented representation for meta learning. In this stacking meta-learning framework, the meta-learner was trained to integrate complementary information from both the peptide descriptors and the prediction patterns generated by heterogeneous base classifiers. After evaluating candidate meta-learners, KNN was selected as the final meta-learner to perform AMP prediction. The predicted probability is computed as the fraction of positive samples among the k nearest neighbors, and the final class label is obtained by thresholding this probability for binary classification.

### Benchmarking with the state-of-the-art methods based on independent datasets

We compared the performance of SAMP V2 with the state-of-the-art AMP prediction methods, including SAMP ^35^, AmPEPpy ^38^, AMPScanner V2 ^39^, AMPpred-MFA ^32^ and iAMP-Attenpred ^30^, as well as five widely used baseline classifiers: SVM ^44^, extreme gradient boosting (XGBoost) ^45^, logistic regression (LR) ^46^, k-nearest neighbor (KNN) ^47^, least absolute shrinkage and selection operator (LASSO) ^48^. To ensure a fair comparison, all models were trained using the same training dataset and evaluated on an identical independent test dataset.

To evaluate the final classification results and facilitate comparison with state-of-the-art models, four commonly adopted performance metrics were calculated, including accuracy, F1-score, MCC, and G-measure. Accuracy measures the overall proportion of correctly classified samples, whereas F1-score provides a balanced assessment by jointly considering precision and recall. MCC offers a comprehensive evaluation by accounting for all four components of the confusion matrix and is particularly suitable for imbalanced classification tasks such as AMP prediction. In addition, G-measure quantifies the balance between correctly identifying AMPs (true positives) and correctly rejecting non-AMPs (true negatives), thereby reflecting performance on both classes and mitigating the bias of accuracy under class imbalance. Together, these metrics provide a comprehensive and unbiased evaluation of model performance on imbalanced AMP datasets. The four evaluation metrics are defined as follows:

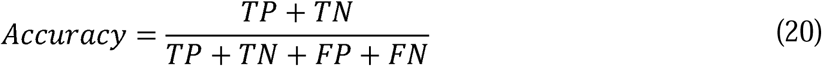

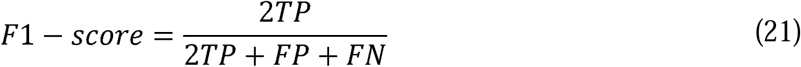

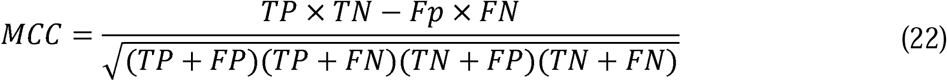

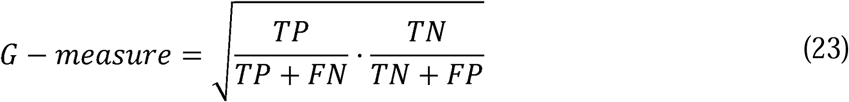

where TP, TN, FP, and FN denote true positives, true negatives, false positives, and false negatives, respectively.

## Results

### Impact of peptide features on predictive performance

To investigate the contribution of different feature types to the stacking model, three categories of sequence descriptors for each peptide were extracted, i.e., AAC, PHYC and BIA-SAAC. We further constructed pairwise combinations (AAC+PHYC, AAC+STRL, PHYC+STRL) and the full combination (AAC+PHYC+STRL). Models trained with each feature set were evaluated on multiple independent test datasets (benchmark1, benchmark2, amphibian, bacteria, human, and plant dataset) using four metrices (Accuracy, MCC, F1-score, and G-measure), with the best-performing configuration per dataset highlighted in bold (**Supplementary Table 1**).

Overall, AAC+STRL showed strong performance across majority independent test datasets, achieving the best or near-best results for all metrics. In contrast, models trained with PHYC alone yielded substantially weaker performance, indicating limited discriminative power when physicochemical descriptors are used in isolation. Importantly, STRL alone already provided competitive performance relative to AAC and other combinations, demonstrating that augmented split amino acid composition captures informative sequence patterns relevant to AMP identification. However, STRL was not uniformly optimal across majority datasets and all metrics, whereas combining it with AAC produced the most stable and consistently improved performance. These results highlight the importance of STRL as an informative feature family and further suggest that its complementary information to AAC is effectively exploited by the stacking framework to improve generalization on independent datasets.

Additionally, the influence of the BIA-SAAC hyperparameter lambda (the maximum sequence-order correlation tier) on model performance was investigated. We evaluated lambda = 1/3/5 on six datasets (**Supplementary Fig. S2**). Overall, increasing lambda led to consistent or modestly improved performance across datasets, with lambda = 5 achieving the best results in most cases. In particular, lambda = 5 provided the highest overall performance for the bacteria, human, and benchmark1 datasets. For the amphibian dataset, lambda = 5 produced the highest accuracy (0.94) and a competitive MCC (0.88). For the plant dataset, lambda = 1 achieved marginally better F1 (0.63) and G-measure (0.89), whereas lambda = 5 remained comparable across metrics. Benchmark1 showed better classification performance with lambda = 5 giving a slightly higher accuracy (0.75), MCC (0.50) and F1-score (0.73). Based on these results, we selected lambda=5 as the default setting for SAMP V2, as it provides the most robust and consistently strong performance across diverse datasets.

### Characterization of the sequence-based features of AMPs

Previous studies have suggested that different regions of peptide sequences may contain region-specific residue patterns that are associated with biological function. In particular, short sequence segments around the N-terminal and C-terminal regions can encode important compositional or sorting-related information, while the middle region may also contribute sequence-order and physicochemical patterns relevant to peptide activity. Therefore, in addition to global amino acid composition (AAC), we further characterized augmented split amino acid composition (BIA-SAAC), which captures both regional residue composition and biochemical-sequence-order correlations from the N-terminal, middle, and C-terminal regions.

The distribution pattern of both AAC and BIA-SAAC were compared between AMP and non-AMP peptides across seven datasets. AMP and non-AMP peptides exhibited distinct global AAC profiles (**Fig. 2A**). Several residues showed clear compositional differences between the two groups. The largest group differences were observed for cysteine (C), lysine (K), glutamic acid (E), glycine (G), aspartic acid (D), and serine (S), marking them as the most differentially represented residues in this analysis. AMP sequences were relatively enriched in residues such as cysteine (C) lysine (K) and glycine (G), whereas non-AMP sequences showed higher frequencies of several other residues, including glutamic acid (E), aspartic acid (D) and serine (S). These compositional differences are consistent with the known cationic and amphipathic properties of many AMPs.

**Fig. 2.**
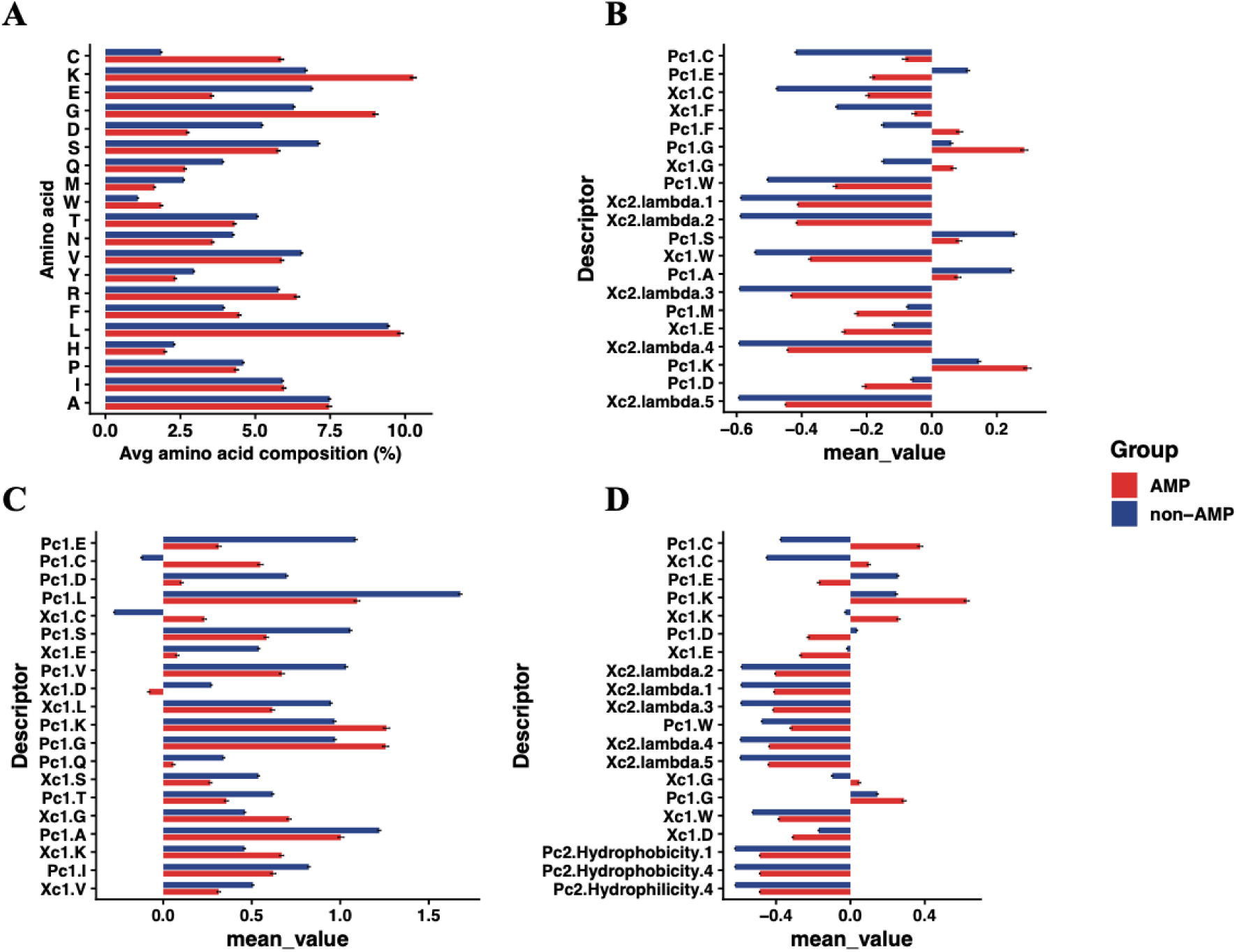
Comparisons of AAC and BIA-SAAC between the AMP and non-AMP peptides based on seven datasets. The frequency of 20 common amino acids within seven datasets were calculated to represent the difference between AMP and non-AMP sequences (**A**). Amino acids are ordered by the absolute difference in mean composition between AMP and non-AMP groups. Biochemical-information augmented split amino acid composition (BIA-SAAC) descriptors were calculated separately for the N-terminal (**B**), middle (**C**), and C-terminal (**D**) peptide regions. For each region, descriptors are ranked by the absolute difference in mean descriptor value between AMP and non-AMP groups. Error bars indicate standard error of the mean. Xc1 descriptors represent PseAAC composition components for individual amino acids, whereas Xc2 descriptors represent PseAAC sequence-order correlation components. Pc1 descriptors represent APAAC composition components, and Pc2 descriptors represent APAAC sequence-order correlation components based on hydrophobicity and hydrophilicity. For example, Xc1.L denotes the PseAAC composition component for leucine, and Xc2.lambda.2 denotes the second-tier sequence-order correlation factor.

We next examined whether regional BIA-SAAC descriptors could provide additional information beyond global AAC. For each peptide, BIA-SAAC features were calculated separately from the N-terminal, middle, and C-terminal regions. AMP and non-AMP peptides displayed distinct regional BIA-SAAC patterns in all three segments (**Fig. 2B-D**). The top-ranked descriptors included both composition-like components, such as Xc1 and Pc1 descriptors for individual amino acids, and sequence-order correlation components, such as Xc2.lambda and Pc2 hydrophobicity/hydrophilicity descriptors. In the N-terminal region (**Fig. 2B**), the highest-ranked descriptors included both amino acid composition-related terms and sequence-order correlation terms, suggesting that N-terminal differences between AMP and non-AMP peptides involve both residue usage and local sequence-order properties. In the middle region (**Fig. 2C**), several high-ranking descriptors were dominated by Pc1 and Xc1 composition components, indicating that regional amino acid composition contributed strongly to group separation. In the C-terminal region (**Fig. 2D**), top-ranked descriptors included cysteine-, glutamic acid-, and lysine-related composition terms, together with Xc2 and Pc2 sequence-order correlation descriptors, highlighting region-specific differences in both composition and physicochemical sequence-order patterns. Importantly, the most discriminative descriptors differed among the N-terminal, middle, and C-terminal regions, suggesting AMP-associated sequence patterns are region dependent rather than uniformly distributed across the peptide sequence. These observations support the use of AAC and BIA-SAAC as complementary sequence-based features for AMP identification in SAMP V2.

### Model Performance and benchmarking with the state-of-the-art methods

Benchmark evaluation on six independent test datasets (amphibian, bacteria, human, plant, benchmark1 and benchmark2) was performed. For each dataset, performance of SAMP V2 was compared against five base classifiers where SAMP V2 consistently achieved the best performance across all six independent datasets (**Fig. 3**), exhibiting the highest values for accuracy, MCC, F1-score, and G-measure. In contrast, the base classifiers showed noticeably lower performance and greater variability in ranking across datasets, indicating that their generalization ability is more sensitive to dataset shifts. Collectively, these results demonstrate that SAMP V2 provides superior and more robust generalization compared with individual base learners when tested on independent datasets, supporting the advantage of the stacking ensemble for AMP identification.

**Fig. 3.**
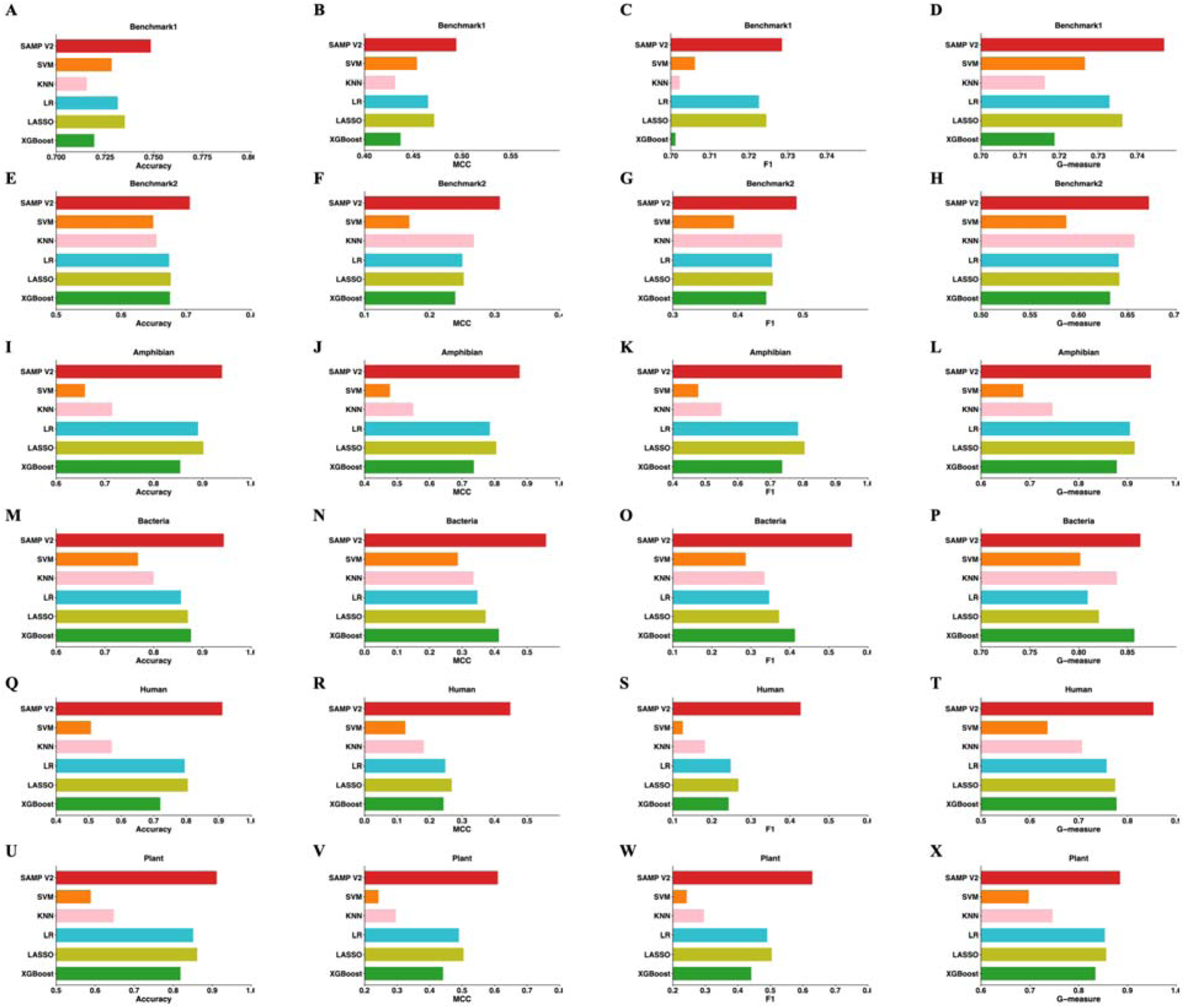
SAMP V2 significantly outperforms the five base classifiers for AMP identification. Comparing SAMP V2 with SVM, XGBoost, LASSO, LR and KNN, in terms of Accuracy (**A, E, I, M, Q, U**), MCC (**B, F, J, N, R, V**), F1-score (**C, G, K, O, S, W**) and G-measure (**D, H, L, P, T,x)**across amphibian, bacteria, human, plant, benchmark1 and benchmark2 dataset, respectively.

To demonstrate the superiority of SAMP V2, we also benchmarked it against state-of-the-art AMP prediction tools, including SAMP, AmPEPpy, AMPScanner V2, AMPpred-MFA and iAMP-Attenpred. These methods represent diverse modeling paradigms, ranging from ensemble learning with random projection (SAMP), conventional RF-based classifiers using physicochemical descriptors (AmPEPpy), deep learning architectures incorporating convolutional, pooling, and recurrent layers (AMPScanner V2), to attention-enhanced frameworks that integrate contextual sequence representations or multi-feature embeddings for AMP classification (iAMP-Attenpred and AMPpred-MFA). Overall, SAMP V2 consistently achieved superior performance across all evaluated metrics (**Figs. 4-5**). Notably, SAMP V2 outperformed both traditional ML classifiers and DL-based approaches, indicating that its ensemble stacking framework effectively integrates complementary information captured by heterogeneous base learners. This advantage was particularly evident in MCC, which are more sensitive to class imbalance and better reflect prediction robustness in AMP identification tasks. These results suggest that stacking-based ensemble learning provides a more effective strategy for AMP prediction than single-model approaches or homogeneous ensembles, by leveraging diverse decision boundaries and reducing model-specific biases.

**Fig. 4.**
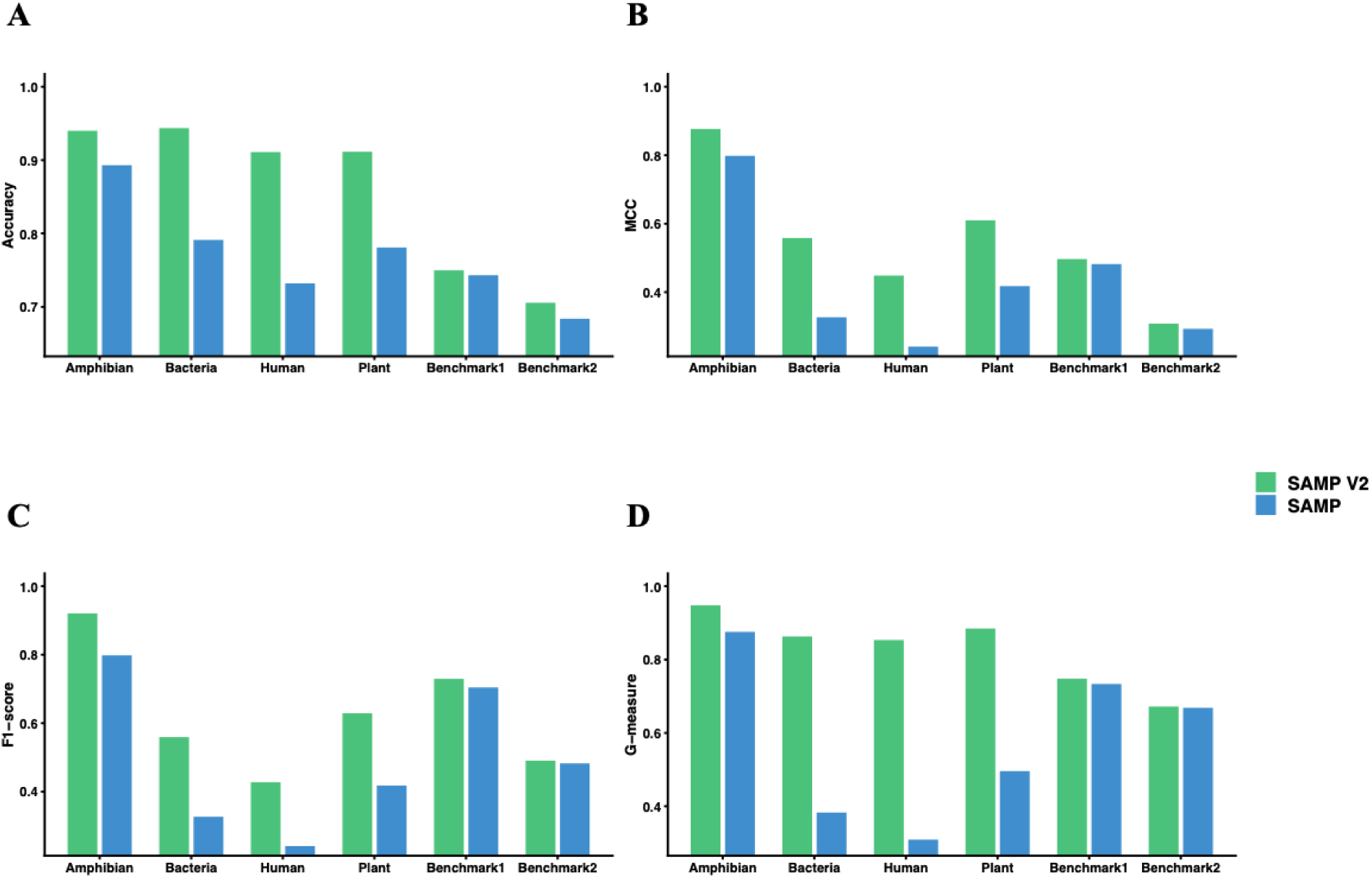
Performance comparison between SAMP and SAMP V2 across benchmark datasets. Comparison of SAMP and SAMP V2 performance across six evaluation datasets, including two benchmark1, benchmark2, amphibian, bacteria, human, and plant. Four performance metrics were evaluated: accuracy (**A**), MCC (**B**), F1-score (**C**), and G-measure (**D**). Green bars represent SAMP V2 and blue bars represent SAMP.

**Fig. 5.**
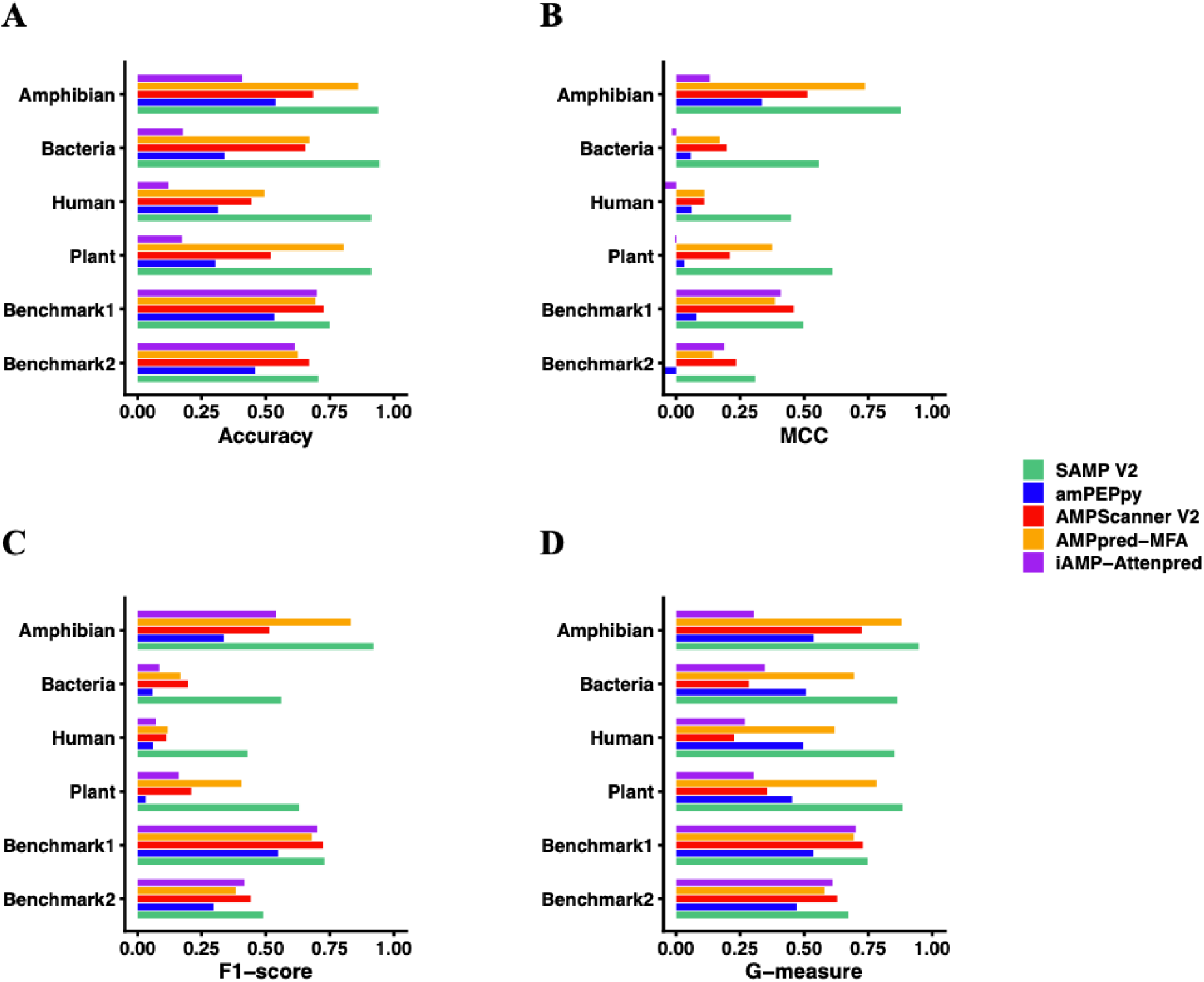
SAMP V2 significantly outperforms the state-of-the-art methods for AMP identification. Comparing SAMP V2 with amPEPpy, AMPScanner V2, AMPpred-MFA and iAMP-Attenpred in terms of Accuracy (**A**), MCC (**B**), F1-score (**C**) and G-measure (**D**) across amphibian, bacteria, human, plant, benchmark1 and benchmark2 dataset. SAMP V2 is shown in red, amPEPpy in green, and AMPScanner v2 in orange.

## Discussion

In this study, we developed SAMP V2, an enhanced computational framework for AMP identification that couples a stacking ensemble learning architecture with a newly proposed sequence representation, BIA-SAAC. Rather than relying on a single classifier or purely global composition features, SAMP V2 integrates complementary decision patterns learned by heterogeneous base models through a meta-learning stage, yielding a more robust and generalizable predictor. In parallel, BIA-SAAC provides a biologically informed encoding that captures both regional heterogeneity across peptide terminals and the biochemical-sequence-order correlations that are closely related to amphiphilicity and structural organization of AMPs. Together, the stacking strategy and the BIA-SAAC representation enable SAMP V2 to better characterize function-relevant sequence patterns and mitigate model-specific bias, thereby strengthening AMP identification under diverse data conditions.

Traditional feature encoding methods, such as AAC, reduce a protein sequence to a fixed-length vector of residue frequencies. While computationally efficient, AAC completely discards sequence-order information. Peptides with the same length would have the exact same AAC vector. PAAC advanced the field by incorporating sequence-order correlations through physicochemical tiers, blending the global composition with correlation factors derived from properties like hydrophobicity and hydrophilicity ^37^. However, even standard PAAC typically operates globally. It calculates correlations across the entire chain and averages physicochemical properties over the full length of the molecule. This global averaging assumes that the biological function is isotropically distributed that a hydrophobic residue at the N-terminus has the same functional implication as a hydrophobic residue at the C-terminus. For cytosolic globular proteins, this approximation might hold some validity. For membrane-active peptides, however, this assumption is fundamentally flawed ^49^. AMPs are structurally and functionally polarized molecules. Their activity depends not just on what amino acids are present, but where they are located relative to the membrane interface ^6^.

In this study, we designed BIA-SAAC to better reflect the fact that AMP activity is often governed by region-specific sequence functions rather than uniform properties averaged over the entire chain. Classical mechanistic models of AMP action, such as the carpet, barrel-stave, and toroidal pore models, share a broadly similar sequence of events, including electrostatic attraction, interfacial partitioning, structural rearrangement, and membrane insertion ^50^. Importantly, these stages are frequently decoupled along the peptide backbone, meaning that different segments of the same peptide can contribute disproportionately to distinct steps of membrane engagement and disruption. This provides a biological rationale for representing a peptide as multiple segments instead of a single global feature vector. By computing pseudo amino acid composition within proportionally defined N-terminal, middle, and C-terminal regions, BIA-SAAC is intended to preserve both local sequence-order correlations and segment-specific physicochemical signals that may otherwise be diluted by whole-sequence averaging. The N-terminus of many α-helical AMPs frequently initiates membrane interaction and insertion and is often enriched in hydrophobic residues ^51^. For example, melittin shows strong polarity with a hydrophobic N-terminal domain and a cationic C-terminal domain, and experiments indicate that blocking the N-terminus reduces antimicrobial/hemolytic activity ^52,53^. In contrast, the C-terminus commonly contributes to electrostatic anchoring and target specificity, as bacterial membranes are negatively charged and many AMPs concentrate Lys/Arg near terminal regions ^54^. C-terminal phenomena such as amidation can further increase net positive charge and stabilize helical structure, highlighting localized physicochemical signatures that global features may obscure.

A further design principle in BIA-SAAC is the augmented splitting scheme, which addresses the substantial length heterogeneity of AMPs. Biological sequences span a wide range of lengths, and for short peptides, a fixed N-terminal window may encompass most of the sequence, leaving negligible middle and C-terminal regions. For longer peptides, the same window captures only a small fraction of the N-terminal helix and may miss functionally relevant positions. By contrast, proportional splitting normalizes segment boundaries across peptides and helps align functional phases (initiation, propagation, termination) across variable-length sequences. As a result, “Segment 1” consistently corresponds to an initiation/insertion domain and “Segment 3” to a terminal anchoring domain regardless of whether peptide residues are long or short, improving comparability across datasets.

SAMP V2 also benefits substantially from its stacking ensemble design. In SAMP V2, heterogeneous base classifiers (SVM and KNN) are trained in parallel to learn complementary decision patterns from the same peptide feature space, and their prediction scores are then integrated by a sparse meta-learner (KNN) together with the original feature representations. This stacking strategy allows SAMP V2 to exploit the complementary strengths of different model families, margin-based separation from SVM and neighborhood-based decision making from KNN, while the meta-learner adaptively assigns weights to the most informative signals and suppresses redundant or noisy ones. As a result, the final stacked predictor can effectively learn from both the raw feature space and the diverse base-model outputs, improving robustness and generalization beyond what can be achieved by any single classifier alone. Similarly, our benchmarking results indicate that SAMP V2 achieves more stable and accurate performance across multiple evaluation metrics compared with individual baseline models, highlighting the advantage of stacking for AMP identification under heterogeneous sequence characteristics and class-imbalance settings.

SAMP V2 demonstrated superior performance compared with existing state-of-the-art AMP identification methods. We benchmarked SAMP V2 against established tools, including SAMP, AmPEPpy, AMPScanner V2, iAMP-attenpred and AMPpred-MFA, as well as five widely used ML classifiers (SVM, LASSO, XGBoost, LR, KNN) trained under the same data setting. Across these evaluations, SAMP V2 consistently achieved stronger performance on multiple metrics, indicating improved robustness and generalization. A possible explanation for this advantage is that SAMP V2 integrates two key innovations: the proposed BIA-SAAC representation, which captures region-specific sequence heterogeneity together with sequence-order correlations that are diluted in global descriptors, and a stacking ensemble learning strategy that combines complementary decision patterns from heterogeneous base learners through a KNN meta-learner. Unlike approaches that rely on a single model type or simple score averaging, the stacking framework in SAMP V2 adaptively learns how to weight base-model outputs while leveraging the original feature space, thereby yielding more accurate and stable AMP predictions.

## Conclusion

In this study, we developed SAMP V2, a stacking ensemble learning framework based on biochemical information and sequence-order information augmented split amino acid composition (BIA-SAAC) for accurate AMP identification. BIA-SAAC extends SAMP by integrating regional amino acid composition with biochemical properties and sequence-order information, enabling a more comprehensive characterization of AMP sequences. SAMP V2 combines complementary predictions from heterogeneous base learners through a KNN-based meta-learner. Across benchmark evaluations, SAMP V2 achieved improved performance in terms of accuracy, F1-score, MCC, and G-measure compared with conventional ML classifiers and state-of-the-art AMP predictors, including SAMP, amPEPpy, AMPScanner V2, iAMP-attenpred and AMPpred-MFA. Overall, SAMP V2 offers a powerful framework for high-throughput AMP screening and prioritization of promising AMP candidates against multidrug-resistant pathogens.

## Supporting Information

**Supplementary Fig. S1 Overview of the datasets used in this study for AMP classification.** The composition of peptide sequence for AMPs (**A**) and non-AMPs (**B**) across datasets. The density distribution of sequence length for AMPs (**C**) and non-AMPs (**D**) across datasets.

**Supplementary Fig. S2 Sensitivity analysis of the BIA-SAAC hyperparameter across six independent test datasets**. Performance comparison of SAMP V2 under lambda =1/3/5 on amphibian (**A**), bacteria (**B**), human (**C**), plant (**D**), benchmark1 (**E**), and benchmark2 (**F**), evaluated by accuracy, MCC, F1-score, and G-measure.

**Supplementary Table 1 Performance comparison of different feature-based models**. To find the best combination of feature types, all combination of different feature categories were tested, including AAC, PHYC, STRL, AAC+PHYC, AAC+STRL, PHYC+STRL, and AAC+PHYC+STRL. The performance of SMAP V2 on six independent test datasets with each feature combination was evaluated. Numbers in bold represent the best performance for each feature scheme.

## Conflict of Interest

The authors declare no competing interests.

## Funding

Research reported in this publication was supported by the U.S. National Science Foundation under Award Numbers 2500836 and 2614824, the National Institute Of Alcohol Abuse And Alcoholism of the National Institutes of Health under Award Number R21AA032098, and the Office Of The Director, National Institutes Of Health of the National Institutes of Health under Award Number R03OD038391. This research was supported by the State of Nebraska through the Pediatric Cancer Research Group, part of the Child Health Research Institute. This work was also partially supported by the University of Nebraska Collaboration Initiative Grant from the Nebraska Research Initiative (NRI). The content is solely the responsibility of the authors and does not necessarily represent the official views of the funding organizations.

## Author contributions

SW conceived and designed the study. SW and MS developed the method, performed the experiments and analyzed the data. All authors participated in writing and revising the paper. The manuscript was approved by all authors.

## Data accessibility

All the data used in this manuscript are publicly available in the corresponding references. SAMP V2 package is available in GitHub at https://github.com/wan-mlab/SAMP-V2.

## Notes

### Competing Interest Statement

The authors have declared no competing interest.

