## Supplementary figures and images for "SAMP V2: A novel stacking ensemble learning model for antimicrobial peptides identification based on augmented split amino acid composition with biochemical-sequence-order information"

### Supplementary Fig. S1

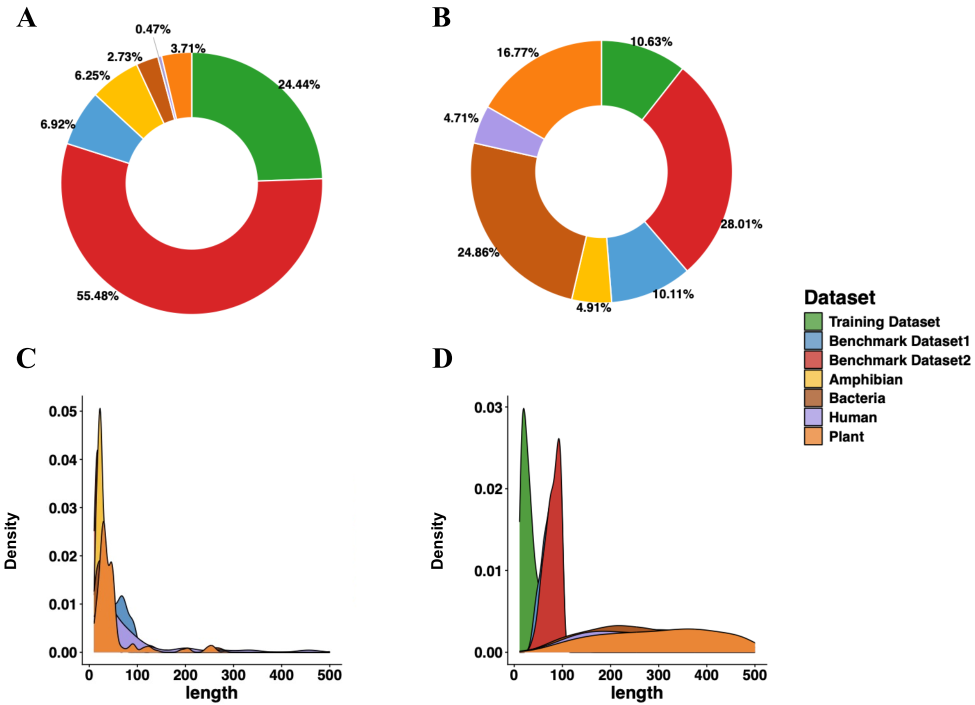

### Supplementary Fig. S2

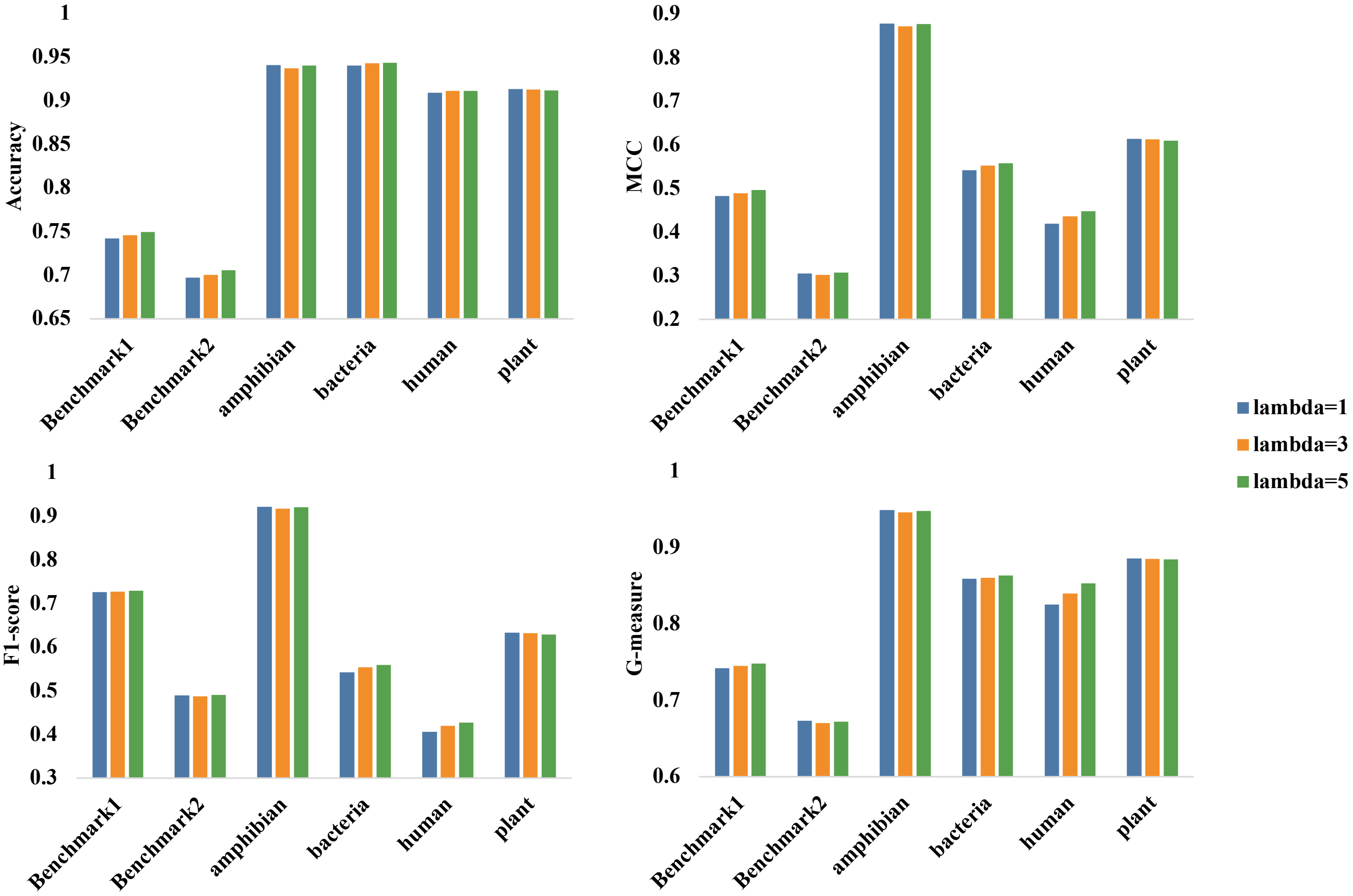
