## Supplemental Table 1 for "SAMP V2: A novel stacking ensemble learning model for antimicrobial peptides identification based on augmented split amino acid composition with biochemical-sequence-order information"

**Supplementary Table 1 Performance comparison of different feature-based models**

| **Dataset** | **Model** | **Acc** | **MCC** | **F1-score** | **G-measure** |
| --- | --- | --- | --- | --- | --- |
| Amphibian | AAC | 93 | 86 | 91 | 94 |
|  | PHYC | 60 | 18 | 50 | 59 |
|  | STRL | **94** | **88** | **92** | **95** |
|  | AAC+PHYC | 76 | 58 | 74 | 79 |
|  | AAC+STRL | **94** | **88** | **92** | **95** |
|  | PHYC+STRL | 93 | 85 | 90 | 94 |
|  | AAC+PHYC+STRL | 93 | 85 | 91 | 94 |
| Bacteria | AAC | 91 | 46 | 45 | 85 |
|  | PHYC | 66 | 5 | 11 | 55 |
|  | STRL | **94** | 53 | 53 | **86** |
|  | AAC+PHYC | 69 | 15 | 16 | 67 |
|  | AAC+STRL | **94** | **54** | **54** | **86** |
|  | PHYC+STRL | 93 | 51 | 50 | 86 |
|  | AAC+PHYC+STRL | 93 | 51 | 50 | 86 |
| Human | AAC | 90 | 40 | 39 | 82 |
|  | PHYC | 50 | 11 | 12 | 62 |
|  | STRL | **91** | **42** | **41** | **83** |
|  | AAC+PHYC | 55 | 12 | 12 | 64 |
|  | AAC+STRL | **91** | **42** | **41** | **83** |
|  | PHYC+STRL | 86 | 34 | 31 | 81 |
|  | AAC+PHYC+STRL | 86 | 35 | 31 | 83 |
| Plant | AAC | 89 | 57 | 58 | 87 |
|  | PHYC | 57 | 6 | 18 | 55 |
|  | STRL | **91** | **61** | **63** | 88 |
|  | AAC+PHYC | 62 | 11 | 21 | 60 |
|  | AAC+STRL | **91** | **61** | **63** | **89** |
|  | PHYC+STRL | 89 | 55 | 57 | 86 |
|  | AAC+PHYC+STRL | 89 | 55 | 57 | 86 |
| Benchmark1 | AAC | 71 | 42 | 70 | 71 |
|  | PHYC | 58 | 17 | 58 | 58 |
|  | STRL | 74 | 48 | 72 | 74 |
|  | AAC+PHYC | 59 | 19 | 59 | 60 |
|  | AAC+STRL | 74 | 48 | 73 | 74 |
|  | PHYC+STRL | 74 | 49 | **74** | **76** |
|  | AAC+PHYC+STRL | **75** | **50** | **74** | 75 |
| Benchmark2 | AAC | 66 | 25 | 46 | 64 |
|  | PHYC | 56 | 3 | 31 | 51 |
|  | STRL | 69 | 30 | 48 | 67 |
|  | AAC+PHYC | 57 | 4 | 32 | 51 |
|  | AAC+STRL | **70** | **31** | **49** | 67 |
|  | PHYC+STRL | 69 | 30 | **49** | 67 |
|  | AAC+PHYC+STRL | **70** | **31** | **49** | **68** |

Acc, accuracy; MCC, Matthews correlation coefficient; G-measure, the geometric mean of recall and precision. Numbers in bold represent the best performance for each test dataset.
